# DNA barcodes and reliable molecular identifications in a diverse group of invasive pests: lessons from *Bactrocera* fruit flies on variation across the COI gene, introgression, and standardization

**DOI:** 10.1101/2020.11.23.394510

**Authors:** Camiel Doorenweerd, Michael San Jose, Luc Leblanc, Norman Barr, Scott Geib, Arthur Y.C. Chung, Julian Dupuis, Arni Ekayanti, Elaida R. Fiegalan, K.S. Hemachandra, M. Aftab Hossain, Chia-Lung Huang, Yu-Feng Hsu, Kimberly Y. Morris, Andi Maryani Mustapeng, Jerome Niogret, Thai Hong Pham, U.G.A.I. Sirisena, Daniel Rubinoff

**Affiliations:** University of Hawaii, College of Tropical Agriculture and Human Resources, Department of Plant and Environmental Protection Sciences, Entomology section, 3050 Maile Way, Honolulu, Hawaii, 96822-2231, USA; University of Idaho, Department of Entomology, Plant Pathology and Nematology, 875 Perimeter Drive, MS2329, Moscow, Idaho, 83844-2329, United States; United States Department of Agriculture, Animal and Plant Health Inspection Service, Plant Protection and Quarantine, Science & Technology, Mission Laboratory, 22675 N. Moorefield Road, Edinburg, Texas 78541, USA; Tropical Crop and Commodity Protection Research Unit, Daniel K Inouye U.S. Pacific Basin Agricultural Center, USDA Agricultural Research Services, Hilo, Hawaii, USA; Forest Research Centre, Forestry Department, P.O. Box 1407, 90715 Sandakan, Sabah, Malaysia; University of Kentucky, Department of Entomology, S-225 Ag Science Center North, 1100 South Limestone, Lexington, Kentucky, 40546-0091, USA; Niogret Ecology Consulting LLC, Wotu, Luwu Timor, Sulawesi Seleaton 92971, Indonesia; Department of Crop Protection, College of Agriculture, Central Luzon State University, Science City of Muñoz, Nueva Ecija 3119, Philippines; Faculty of Agriculture, University of Peradeniya, Sri Lanka; Insect Biotechnology Division, Institute of Flood and Radiation Biology, Bangladesh, Atomic Energy Commission, Dhaka-1349, Bangladesh; Institute of Oceanography, Minjiang University, No. 200, Xiyuangong Road, Shangije Town, Minhou County, Fuzhou City, Fujian Province, China; Department of Life Science, National Taiwan Normal University, No. 88, Sec. 4, Tingzhou Road, Taipei, 116, Taiwan, ROC; Mientrung Institute for Scientific Research, Vietnam Academy of Science and Technology, 321 Huynh Thuc Khang, Hue, Vietnam; Vietnam National Museum of Nature & Graduate School of Science and Technology, Hanoi, Vietnam; Department of Plant Sciences, Faculty of Agriculture, Rajarata University of Sri Lanka, Sri Lanka

**Keywords:** Oriental fruit fly, DNA barcoding, Tephritidae, Dacinae, diagnostics, taxonomy

## Abstract

The utility of a universal DNA ‘barcode’ fragment of 658 base pairs of the *Cytochrome C Oxidase I* (COI) gene for the recognition of all animal species has been a widely debated topic on theoretical and practical levels. Regardless of its challenges, large amounts of COI sequence data have been produced in the last two decades. To optimally use the data towards reliable species identification will require further steps to validate the method and reference libraries. The fruit fly tribe Dacini holds about a thousand species, of which eighty are pests of economic concern, including some of the world’s foremost fruit and vegetable pests, and there are many morphologically cryptic species complexes in the tribe. Where previous studies showed limited success in using COI to identify Dacini, our results with a highly curated morphological dataset indicate high congruence between morphology and COI: 90% of the species in our 5,576 sequences, 262-species global dataset can be identified with COI alone based on a monophyly criterion. However, in some key pest species belonging to complexes that were previously thought diagnosable with COI, we found that expanded sampling and independent validation of identifications using genomic data revealed introgression of mitochondrial DNA. We find that the informative SNPs are uniformly distributed across the COI gene, and we provide recommendations for standardization. We conclude that reliable molecular identifications with COI require extensive species coverage, population sampling, and genomics-supported reference identifications before they can be validated as a “diagnostic” marker for specific groups.

## Introduction

The premise and promise of DNA barcoding was that a universal short segment of DNA could be used for species discovery, species delimitation, and to deliver robust species-level identifications without specialized knowledge from taxonomic experts (Hebert et al. 2003b, 2003a). In an era where ‘classic’ taxonomy was no longer viewed as a profitable science – evident from the lack of funding opportunities and quickly vanishing professional taxonomist positions (Sangster and Luksenburg 2015; Wilson 2017) – such a concept was quickly adopted by the biological scientific community and some governments to fill the need for species-level designations and sparked the ‘International Barcode of Life’ initiative (Savolainen et al. 2005; Bonants et al. 2010). This opened up new sources of funding and the DNA barcode database (Ratnasingham and Hebert 2007) now holds over 3.9 million species-level DNA barcodes (Sept. 2020), and over 3,700 papers with “DNA barcode” in the abstract have been published in scientific journals since 2004 (DeSalle and Goldstein 2019). However, the accuracy and reliability of DNA barcoding has been the subject of much debate, addressing both theoretical and practical aspects (e.g. (Rubinoff et al. 2006; Roe and Sperling 2007; Krishna Krishnamurthy and Francis 2012; Collins and Cruickshank 2013). Over time, with mounting evidence on the limitations of DNA barcoding across a variety of organismal life (e.g. (van Nieukerken et al. 2012; Smit et al. 2013; Jiang et al. 2014; Gibbs 2017), the concept has evolved to become a complementary identification tool rather than the replacement multi-tool it was initially portrayed to be, but with unique value in that niche. Nonetheless, even within this revised framing, controversy persists on how the molecular data should be generated, analyzed and interpreted (DeSalle and Goldstein 2019).

The reasons DNA barcoding fails in identifications can be assigned to two broad categories: natural and unnatural causes. Natural causes include biological processes such as incomplete lineage sorting and introgression, which can be difficult to recognize and/or mitigate. Unnatural causes, also referred to as ‘operational error’ include clerical mistakes, pseudogene amplification, alignment error, tree inference method, identification mistakes and inaccurate reference taxonomy (Mutanen et al. 2016), which can be prevented or addressed. Studies that used highly curated datasets, as opposed to a general data-pull from openly accessible databanks, revealed that much of the false DNA barcode-based identifications can be traced back to operational error (Mutanen et al. 2016; Pentinsaari et al. 2020). Accounting for both sources of inaccuracy requires establishing accurate DNA barcode reference libraries based on vetted reference taxonomy (e.g. no over-split or lumped taxa), DNA barcode-independent identifications (e.g. based on morphology or genomics), as well as adequate coverage of intraspecific diversity (populations). All of these protocols are essential for accurate identifications. Unfortunately, DNA barcode-generating initiatives have generally been opportunistic and focus on covering as many species as possible with typically fewer than ten individuals per species (e.g. (Kerr et al. 2007; Smit et al. 2013; Mutanen et al. 2016). Because the genetic diversity within species can vary widely even between closely related lineages and involve thousands of unique haplotypes (Doorenweerd et al. 2019b; Phillips et al. 2019), the practical use of many barcode datasets is limited. DNA barcode-independent identifications of reference specimens are crucial to avoid circular reasoning (DeSalle et al. 2005; Collins and Cruickshank 2013), although the barcode sequence data can be used to instigate re-evaluation of other sets of characters. Attempts at automated molecular species delimitation based on DNA barcode data have been numerous, but, due to aforementioned natural processes, will never be 100% accurate (Tautz et al. 2003; Luo et al. 2018) and are prone to circular reasoning. Instead, an integrative approach using multiple sources of information, including morphology, behavior, ecology, biogeography, and increasingly commonly, phylogenomic datasets are crucial to assign initial species boundaries to establish reference taxonomy for the barcodes, essentially providing them with the background data they need to be reasonably accurate. Currently published DNA barcode datasets typically suffer from flaws in one or many of these categories, preventing accurate assessment of the utility of DNA barcodes for species identification even when all operational error has been excluded. If the vast resources poured into generating millions of DNA barcodes are to reach their optimal utility, systematic procedures for establishing their robustness and baseline accuracy are essential (Floyd et al. 2010; Boykin et al. 2012).

The Dacini fruit flies (Diptera: Tephritidae: Dacinae) are an ideal model group for evaluating the utility of DNA barcodes given our improved understanding of their limitations. Dacini holds almost one thousand described species that are morphologically difficult to distinguish even for specialists, and around eighty representatives are important agricultural pests, making accuracy in identification critical for both quarantine and control measures (Vargas et al. 2015; Doorenweerd et al. 2018; San Jose et al. 2018a). Inaccurate identifications of specimens can set off expensive quarantine and control efforts with severe implications for international trade. Unfortunately, the typical set of morphological characters used to identify Dacini to species is limited, and includes equivocal differences like subtle variation in coloration or wing markings. Overall, Dacini morphology can therefore only provide limited information for classification and identification, especially for the non-expert faced with the need for rapid identification. The taxonomy of the group has only recently begun to stabilize, following integrative approaches and particularly the inclusion of molecular methods (Virgilio et al. 2009; Schutze et al. 2015; San Jose et al. 2018a). Molecular identification would be particularly useful for this group because it would be invaluable for species identification by non-experts for pest-management and international quarantines. Moreover, recent advances in taxonomic knowledge of the group (Schutze et al. 2017) allow for a retrospective comparison with previously published DNA barcode datasets used for Dacini and their respective conclusions on the reliability of molecular identifications.

Too often, DNA barcodes are used without excluding operational error to a reasonable extent, and commonly without any assessment of the incidence of introgression and incomplete lineage sorting, leaving their actual accuracy unvalidated (Mutanen et al. 2016). In this study, we aimed to establish the most complete and thorough DNA barcode reference library for Dacini based on intensive morphological study by taxonomic experts and high biogeographic coverage to exclude operational error as much as possible, with the ultimate goal of assessing DNA barcode reliability for molecular identifications. Specifically, given that accuracy is critical in a group like Dacini with dozens of cryptic invasive pests and international ramifications for incorrect identifications, how much data is needed to verify DNA barcodes, and to what extent can they be applied across a tribe? If a methodology can be established for developing a critical minimum for datasets to verify the utility of DNA barcodes, we can understand their pitfalls and they could be used with greater efficacy.

## Material & Methods

### Taxon sampling and species assignment

Our sampling is based on targeted collecting efforts and careful curation of specimens by world-specialists. We aimed to obtain COI sequences from as many Dacini species as possible and attempted to get representatives from throughout their distribution ranges, through intensive collecting in the period of 2006–2019. Previous studies on DNA barcoding of Dacini fruit flies have mostly been regional (Virgilio et al. 2009; Meeyen et al. 2014; Kunprom and Pramual 2019) or were done prior to major changes in the taxonomy of the group (Jiang et al. 2014). Flies were collected in the field by using the male attractants methyl eugenol and cue lure, and since 2016 also using zingerone (Royer 2015). Typically, methods involved a lure plug in a bucket trap along with a small strip infused with dichlorvos as killing agent.

Traps were suspended ~2m from the ground on a branch with at least 3m between the different lures and left out in the field for 2–5 days, after which the flies were transferred into 95% ethanol and stored at −20°C prior to DNA extraction. As a result of using male lures for collection, almost all sampled specimens are males. No immatures were included because they cannot be identified to species-level based on morphology. Further details on the trapping methodology can be found in Leblanc et al. (Leblanc et al. 2015b). Identifications to species-level were initially done based on morphology, and after sequencing COI, they were re-evaluated if suggested by DNA results. Results incongruent with morphology were verified by repeating PCR, and occasionally DNA extraction, and sequencing, to exclude contamination and clerical error. Assignment to morphological complexes follows that of Drew & Romig (2013) and other leading taxonomic resources. In the special cases of *B. rubigina*, *B. osbeckiae* and *B. melastomatos* we tentatively considered these as different forms of the same species based on high resolution genomic data (Dupuis et al. 2018) but we do not synonymize them here; this group will be covered in further detail in a future publication. Several species in the *B. dorsalis* complex posed additional identification challenges: there is documented morphological overlap between species (Leblanc et al. 2015a) and hybridization and introgression (San Jose et al. 2018b). Therefore, for *B. dorsalis*, *B. occipitalis*, *B. kandiensis*, *B. carambolae*, *B. raiensis* and yet undescribed lineages *B.* “spMalaysia14” and *B.* “spOTU5353” we relied on identifications supported by genotype-by-sequencing (GBS) genomic data (San Jose et al. in prep.). Collecting, depository, and identification data for all specimens are available on BOLD [DOI: xxxxx], the classification follows (Doorenweerd et al. 2018). Some specimens formed monophyletic clusters and were morphologically distinct, and likely represent undescribed species: we assigned them putative species names for the purposes of the present study. We included *Ceratitis capitata* (Tephritidae: Dacinae: Ceratitidini) [21 sequences], *C.* sp. [1 sequence] and *C. cosyra* [2 sequences] as outgroups.

### DNA extraction, amplification, and sequencing

We typically used one leg per specimen for DNA extraction, or in some cases, a full specimen when future genomic studies were anticipated. DNA extraction was performed as previously described in (Garzón-Orduña et al. 2020) and (San Jose et al. 2018a). When available, voucher specimens are deposited in the University of Hawaii Insect Museum (UHIM). PCR amplification targeted a 1,493 base-pair (bp) section of the Cytochrome C Oxidase I (COI) gene with forward primer LCO_mod (5’-3’ NNNNNNNNNNNNNNNNTYTCAACAAATCATAAAGATATTGG; Folmer et al. 1994) and Pat (5’-3’ NNNNNNNNNNNNNNNNTCCAATGCACTAATCTGCCATATTA; Simon 1994) (Figure 1). LCO_mod was modified from the standard LCO primer with a mixed base and modification to better match tephritid mitochondrial COI based off of an alignment of publicly available tephritid mitochondrial genomes. The majority of samples were sequenced using the PacBio Sequel II SMRT platform, following a modification to the protocol outlined in (Garzón-Orduña et al. 2020). Like Garzón-Orduña et al. (2020), we created a dual barcoding scheme with unique 16 bp sequence tags added to the 5’ end of the forward and reverse primers (indicated with “N” above), using the Pacific Biosciences RSII 384 v1 suggested list of tags (a copy is available at Dryad DOI: xxx.xxx). We used barcodes 1–96 on the forward primer and 97–192 on reverse primer to enable multiplexing of ~4,000 samples with anticipated >30x read coverage in a single Sequel II run. The PCR reaction was simplified to include 5 μl NEB Q5 Hot Start High Fidelity Master Mix (New England Biolabs), 1μl of the LCO_mod primer, 3 μl of the PAT primer (both added to a final concentration of 0.4 μM), and 1 μl of DNA for a 10 μl reaction volume. In this modified reaction, we used a high-fidelity polymerase to control for PCR errors, and removed the addition of ThermaStop (Sigma-Aldrich), as we did both the target PCR and indexing in a single PCR reaction. Cycling conditions were an initial denature of 94 °C for 3 minutes, followed by 39 cycles of 94 °C for 30 seconds, 50 °C for 30 seconds and 70 °C for 1 minute, followed by a final extension at 70 °C for 10 minutes. For samples sequenced with PacBio, libraries were prepared by pooling equal volume of each PCR reaction, cleaning up the pool with 0.7X AMPure beads (Beckman Coulter) and processing the sample through the SMRTbell Express Template Prep Kit 2.0 (Pacific Biosciences). Sequencing on a PacBio Sequel platform was outsourced to UC Berkeley (Berkeley, CA, USA). CCS (https://github.com/PacificBiosciences/ccs) was used to produce highly accurate consensus sequences, or, HiFi reads, from raw PacBio data (Garzón-Orduña et al. 2020). HiFi reads were demultiplexed using the Mothur 1.42 command fastq.info (Schloss et al. 2009), and consensus sequences of HiFi reads were generated using the script ccs-consensuser.py (https://github.com/popphylotools/ccs-consensuser). Output files were manually checked for heteroplasmy and stopcodons in Geneious R10 (https://www.geneious.com). A small subset of the samples was sequenced using bidirectional Sanger sequencing, which was outsourced to Eurofins (Louisville, KY, USA). For this subset, sequencing was performed in two parts, using primers LCO_mod and HCO-2198 (5’-3’ TAAACTTCTGGGTGTCCAAARAATCA; Folmer et al. 1994) for the COI-5P segment and HCO-2198rc (reverse complement of HCO-2198) and Pat for the COI-3P segment (Figure 1). Geneious R10 was used to assemble sense and antisense strands and create alignments. We checked the final alignment for insertions and deletions, as well as stop-codons, which indicate the presence of nuclear pseudogenes. The sequences are deposited both in BOLD (DOI:xxx.xxx.xxx) and NCBI GenBank, accessions xxxx-xxxx.

**Figure 1:**
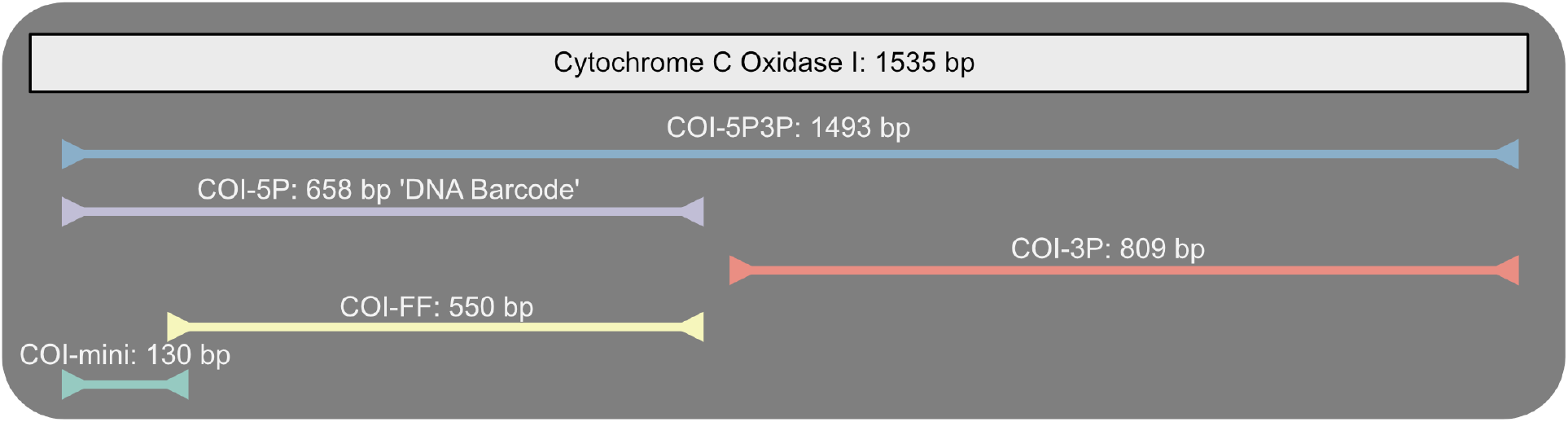
Schematic overview of the Cytochrome C Oxidase I (COI) gene and the segments used for different datasets drawn to scale, bp = base pairs.

### Datasets

Different segments of COI have been proposed and used for species diagnostics, species delimitation and phylogenetics of Dacini (Hebert et al. 2003a; Meusnier et al. 2008; Blacket et al. 2012; Barr et al. 2014; San Jose et al. 2018a). The full COI gene is 1,535 bp long in Dacini (Yong et al. 2016), and because the fragments generated in our study cover 97% of the full gene, we divided the dataset in different segments (Figure 1) and tested for differences in their diagnostic value. We refer to the segment sequenced in our study as COI-5P3P. The COI-5P section is the ‘DNA barcode’ segment (Hebert et al. 2003a), and is prevalent in the Barcode of Life reference database ‘BOLD’ (Ratnasingham and Hebert 2007). The COI-3P segment has been used as one of seven markers in a molecular phylogeny of Dacini (San Jose et al. 2018a), as well as to support species delimitation (Leblanc et al. 2018, 2019) and population source exclusion of *B. dorsalis* (Barr et al. 2014). The COI-FF segment has a forward primer inside the COI-5P fragment to avoid a nuclear pseudogene in the Queensland fruit fly, *Bactrocera tryoni* Frogatt, and is adopted as the standard diagnostic fragment in the Australian Handbook for the Identification of Fruit Flies (Blacket et al. 2012; Schutze et al. 2018). The COI-mini fragment (Meusnier et al. 2008) is the shortest of our set at 130 bp and has potential for use with heavily degraded DNA samples, provided that its discriminatory value is similar to other fragments.

### PyCOIstats and monophyly testing

We created custom Python scripts, the PyCOIstats package (DOI: http://doi.org/10.5281/zenodo.3834896), to analyze our COI data. Most other software that calculates pairwise distances between DNA sequences considers missing or ambiguous data as potential differences, and therefore does not regard sequences with such ‘differences’ as duplicates. However, for the purposes of species identification, we base our statistics on ‘distinct’ haplotypes only (i.e. ambiguities and gaps in otherwise identical sequences do not translate to different haplotypes). Duplicate haplotypes in a dataset can result in underestimating the variability within species and unnecessarily increase computation time. As an exception, the PyCOIstats scripts retain duplicate haplotypes that are shared across species. The escripts can be executed consecutively (i.e. in a ‘pipeline’), depending on the statistics required, and a Jupyter Notebook template is available to plot the data interactively and export graphs for publication. Our first step removed non-unique haplotypes using the q_filter.py script. We then used Geneious R10 to create separate alignments for each of the three Dacini genera and for each COI segment (Figure 1). Pairwise distances were calculated for all alignments with the pdistancer.py script, which calculates intra- and interspecific distances, the maximum intraspecific distance for each species (Dmax) and the minimum distance to the nearest neighbor species (Dmin_NN). The resulting data were plotted with the Jupyter Notebook ‘graphs.ipynb’ using the packages Pandas, Seabourn and MatplotLib. Output files were optimized for publication using Affinity Designer v1.8.3. We used the hapcounter.py script to count the number of unique haplotypes for each species. All analyses were conducted using PyCOIstats v1.3. We used IQ-Tree v1.6.11 (Nguyen et al. 2015) to infer a maximum likelihood tree based on the total dataset with 5,576 aligned COI-5P3P sequences. The IQ-Tree integrated ModelFinder determined TIM2—+F+I+G4 as the best fitting model according to the Bayesian information criterium. The full tree is available in Dryad (DOI:xxx.xxx). To test concordance between COI-based and morphology-based species designations, we used a monophyly criterium (see also (Mutanen et al. 2016); only if the queried taxon is placed in a monophyletic species cluster through phylogenetic tree inference can it be reliably diagnosed. We rooted the IQ-Tree result with the outgroups and used the ‘monophylizer’ perl script (Mutanen et al. 2016) to assess species’ monophyly, polyphyly and paraphyly. The full output table is available in Dryad (DOI:xx.xx).

## Results

The COI reference library we generated contains 5,576 COI-5P3P sequences, of which 3,677 sequences are in the distinct haplotype dataset, representing 262 putative species. The library covers 39 species of *Dacus* [114 distinct sequences], 51 species of *Zeugodacus* [371 distinct sequences] and 163 species of *Bactrocera* [3,172 distinct sequences]. In the total dataset, there are on average 21.3 sequences per species, but there is large variation across the dataset, exemplified by 64 singletons, 52 doubletons, and 2,047 sequences for *B. dorsalis*, mirroring the abundance differences of these flies in different regions (Doorenweerd et al. 2019a).

### Discriminatory power of COI segments

Our results reveal that even though the number of haplotypes increases with longer segment lengths (Fig. 2), the species-discriminatory power is comparable for all segments, except COI-mini (Fig. 3). Thus, although segment COI-5P3P is more than twice as long as all the others, the information from the added genetic variation is countered by increased saturation, and therefore does not result in increased species-level discrimination. Ideally, the genetic variation within species is smaller than the variation between species – i.e. Dmin_NN > Dmax; the so-called barcode-gap – allowing for unambiguous species assignments regardless of using alignment-based distance methods or tree-based monophyly criteria. This is the case for the majority of the species, where Dmax remains below 2-, and Dmin_NN is greater than 2- in *Bactrocera*, greater than 4- in *Zeugodacus*, and greater than 6- in *Dacus* (Fig. 3). However, it should be noted that there are exceptions to these general rules, and no hard p-distance threshold value can be set to accurately delimit all species in our dataset. The only segment that showed a lower discriminatory power is the 130 bp COI-mini, which has a larger spread in Dmax values and overall lower Dmin_NN values, with overlap between both. Thus, although this segment can be used to distinguish some of the species, it is not reliable as a general identification marker.

**Figure 2:**
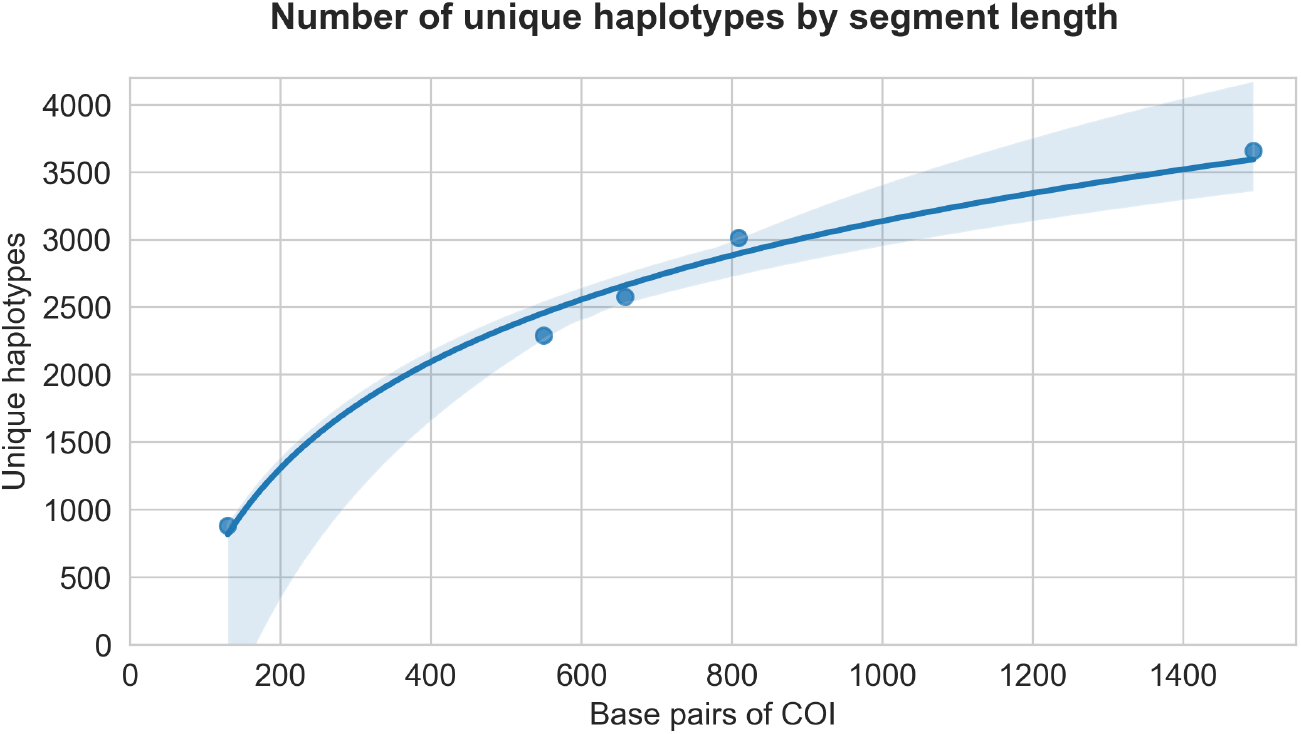
The number of unique haplotypes by segment length for the five datasets (points) based on the same samples for each point, with a log(x) fit of the regression model (line) and shaded confidence interval shows a non-linear increase; indicating increased resolution but also increased saturation at larger segment lengths.

**Figure 3:**
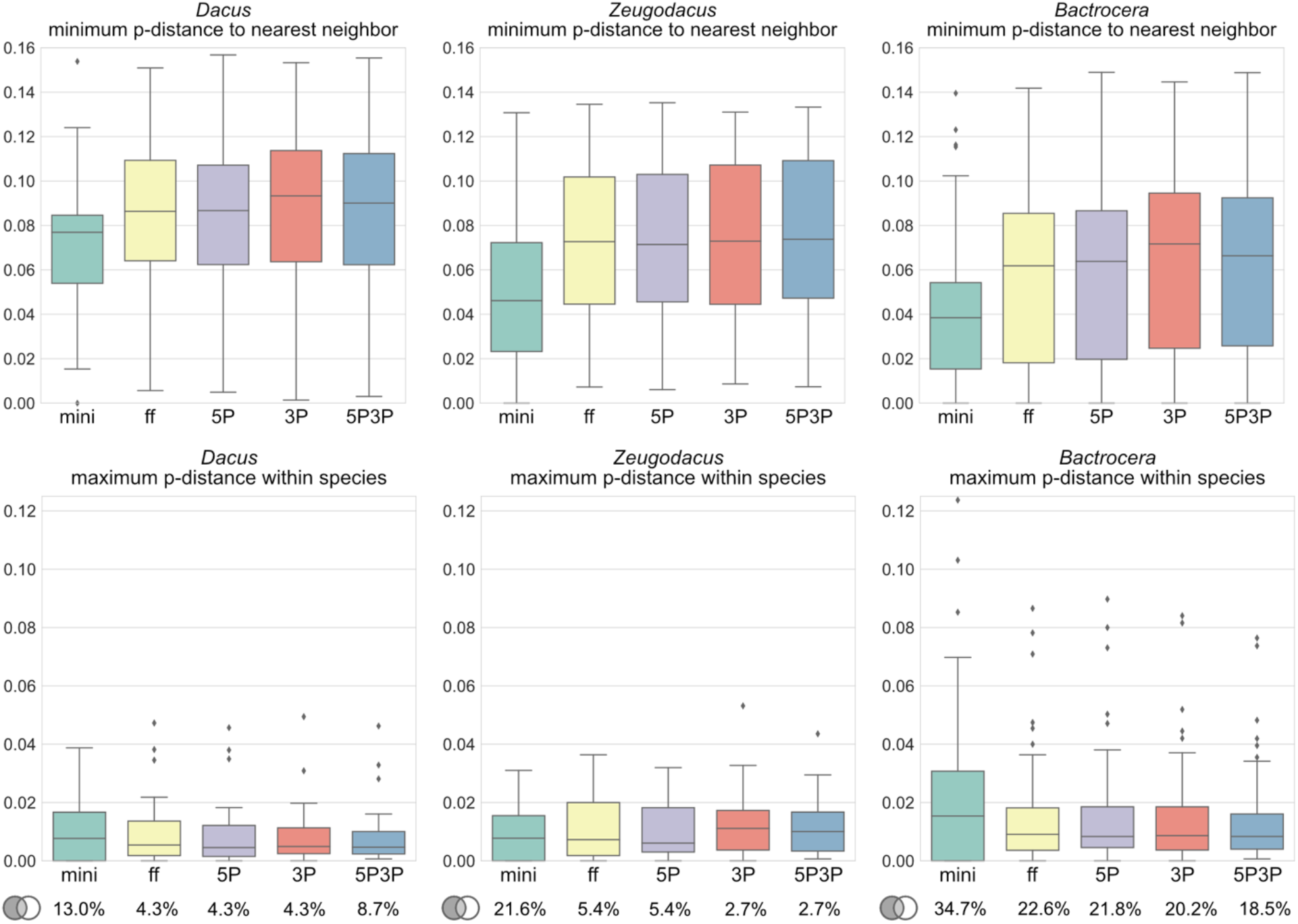
The top row box plots show the minimum p-distance to the nearest neighbor species (Dmin_NN); the bottom row box plots show the maximum p-distance within species (Dmax); across the three genera (columns) based on the different segments of COI. The percentages at the bottom indicate species with Dmax > Dmin_NN (i.e. no barcoding gap) for each genus and segment.

### Differences between genera

We found clear and unexpected differences in relative genetic distances between the three main Dacini genera. The pairwise distance to the nearest neighbor species in *Dacus* is mostly between 6–12%, in *Zeugodacus* it is 4–11% and in *Bactrocera* is 2–10% (Fig. 3); evidencing that COI discriminates best in *Dacus*, followed by *Zeugodacus* and species identification is most challenging in *Bactrocera*. Those differences can be attributed in part to morphological complexes, which are most common in *Bactrocera* (Drew and Romig 2013). However, this would be expected to only affect the lower boundary of Dmin_NN, and not the upper boundary. For *Dacus*, the difference could reflect a sampling artefact, as *Dacus* is most diverse in the Afrotropical region where our sampling coverage was less extensive. However, the distances we find are similar to those in (Virgilio et al. 2012), which was focused on the African continent. A third explanation could involve different evolutionary diversification speeds, and potentially increased speciation levels in *Bactrocera*. The genetic variation we find within species is similar between *Bactrocera* and *Zeugodacus*, but smaller in *Dacus*. However, this could be a sampling artefact as we sampled fewer *Dacus* individuals per species on average; 3.2 specimens per species on average for *Dacus* versus 10.2 for *Zeugodacus* and 29.8 for *Bactrocera* (Supplemental material S2).

### Species monophyly

Based on maximum likelihood (ML) tree inference of COI-5P3P, all included *Zeugodacus* species form monophyletic clades and can thus be identified reliably using COI (summarized results in Table 1; see Dryad DOI:xxxx for the full table and ML tree). Only a single *Dacus* is non-monophyletic. In contrast, 24 species of *Bactrocera* were inferred as non-monophyletic, with most of those part of previously recognized morphological complexes – but the violators of monophyly are not always with members of the same complex. Nine species are part of the *B. dorsalis* complex, four are part of the *B. frauenfeldi* complex, three are part of the *B. nigrotibialis* complex, and four form the *B. tryoni* complex. Only two species are not part of any currently recognized morphological complex. Table 1 also lists what we infer as the suspected cause for the non-monophyly, which can be natural causes such as introgression and incomplete lineage sorting, or operational error, such as inaccurate taxonomy. In some cases, we suspect mtDNA ‘swamping’, which can happen when the mitochondrial genome of one species introgresses into another and provides a selective advantage, swamping out and replacing the original mtDNA throughout all populations of that species (e.g., (Dupuis and Sperling 2015).

**Table 1:**
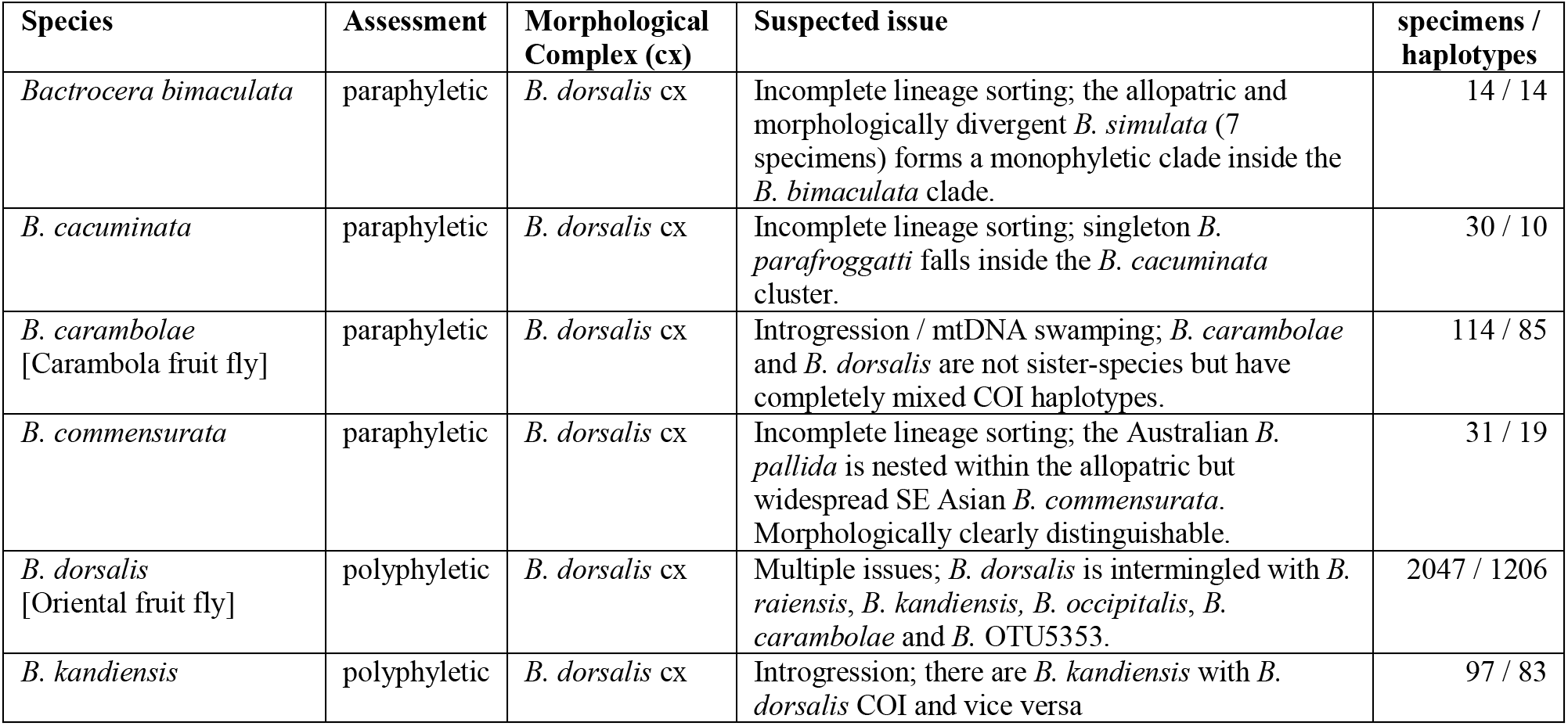
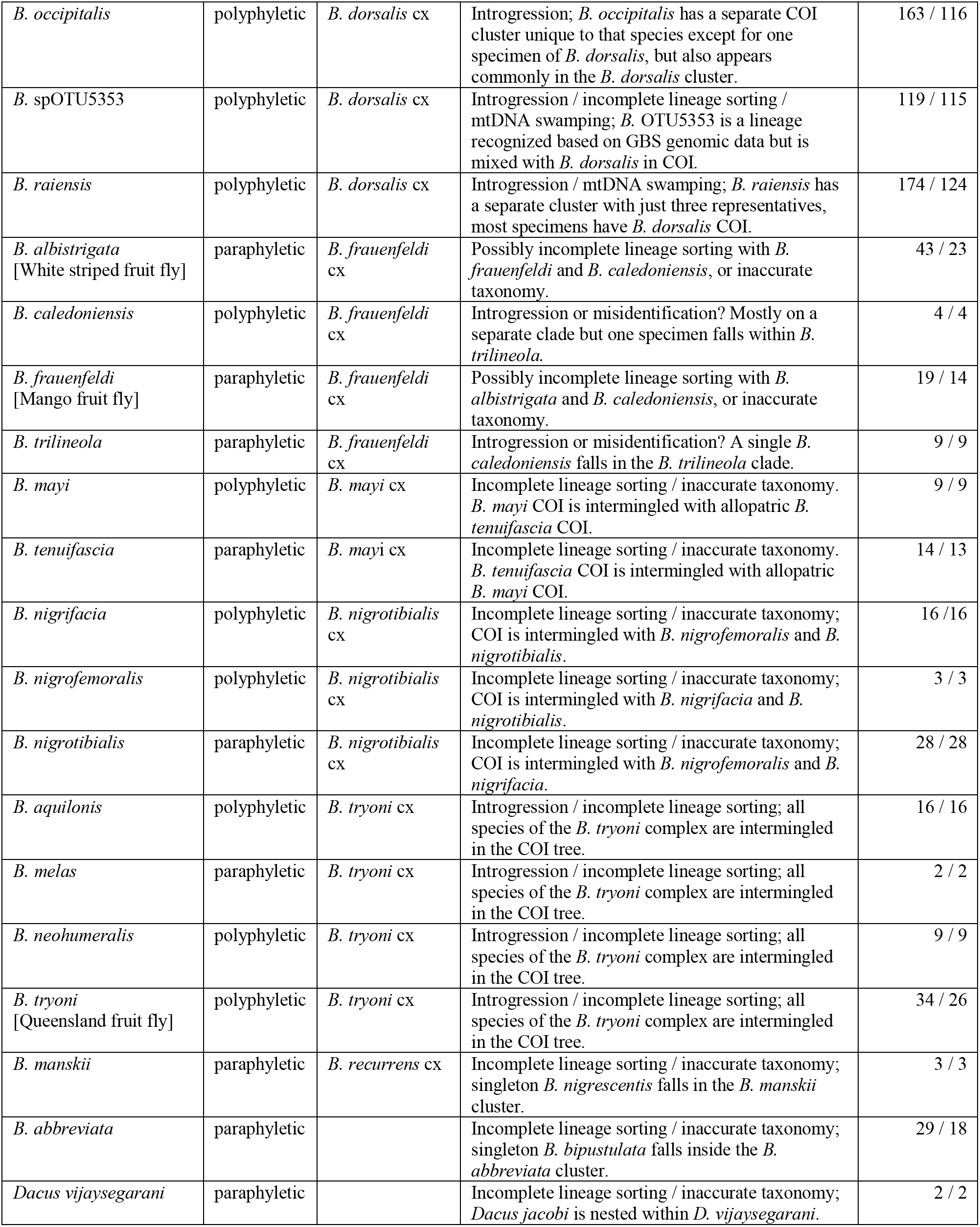
Non-monophyletic species based on maximum likelihood tree inference of COI-5P3P.

*Dacus jacobi* was recently described from India based on morphology (David et al. 2020). We collected specimens in Bangladesh that morphologically match this species, but they form a monophyletic COI cluster within the similar looking *D. vijaysegarani*, which is known from Thailand, Malaysia and Vietnam. The single specimen of the Indian and Sri Lankan *B. bipustulata* that we sampled falls within the COI cluster of the allopatric but widespread Southeast-Asian *B. abbreviata*. *Bactrocera manskii* is known from Queensland and the Torres Straight Is., and in our results its COI is mixed with a singleton *B. nigrescentis*, which is known from Papua and the Solomon Is. Both species are part of the *B. recurrens* morphological complex, and *B. manskii* has been synonymized with *B. recurrens* in the past, but had its status restored based on morphology (Drew 1989). The morphological differences within this complex are minor and species designations appear to rely on geographic distributions. More extensive sampling and study of nuclear DNA can shed light on whether all these are cases of incomplete lineage sorting, introgression or inaccurate taxonomy.

### Bactrocera dorsalis complex

A 10-species monophyletic subset of the 87-species rich *Bactrocera dorsalis* morphological complex, which we refer to as the *B. dorsalis* sensu lato group (San Jose et al. 2018a), has recently been studied with phylogenomic approaches based on genotype-by-sequencing (GBS) single-nucleotide polymorphism (SNP) data as well as 486 single copy ortholog loci, resulting in the first robust phylogeny for this group (San Jose et al. in prep; Fig. 4). We used largely the same set of samples to obtain COI. Using the GBS data to inform species-level identifications where morphology is ambiguous reveals widespread non-monophyly (Table 1). Likely, this is due to introgression or possibly incomplete lineage sorting, summarized in Figure 4. *Bactrocera dorsalis*, Oriental fruit fly, is the most widespread invasive pest of the group and its COI appears to have introgressed into five other species. Vice versa, *B. dorsalis* appears to have received COI DNA from *B.* spOTU5353, *B. occipitalis* and *B. kandiensis*. *Bactrocera kandiensis* associated COI is occasionally recovered in invasive populations of *B. dorsalis* in Africa and Europe (Nugnes et al. 2018). *Bactrocera carambolae* does not have a COI cluster associated only with that species, and it is possible that its COI has completely been replaced with *B. dorsalis* COI. Our data suggest a similar process may be ongoing between *B. raiensis* and *B. dorsalis*; of the 174 *B. raiensis* specimens sampled, only three had COI that did not fall in the *B. dorsalis* cluster but formed a separate cluster that likely represents its original COI. *Bactrocera commensurata* is non-monophyletic with regards to *B. pallida*, which is monophyletic within it, and the pair can be considered as an identification unit for COI. The Pacific *Bactrocera simulata* forms a COI cluster with a monophyletic cluster of the Southeast Asian *B. bimaculata* inside. The latter is part of the morphologic *B. bryoniae* morphological complex; a possible case of incomplete lineage sorting. Similarly, the Pacific singleton *B. parafroggatti* falls within the Australian *B. cacuminata* cluster, but *B. parafroggatti* is morphologically distinct and not part of the *B. dorsalis* complex.

**Figure 4:**
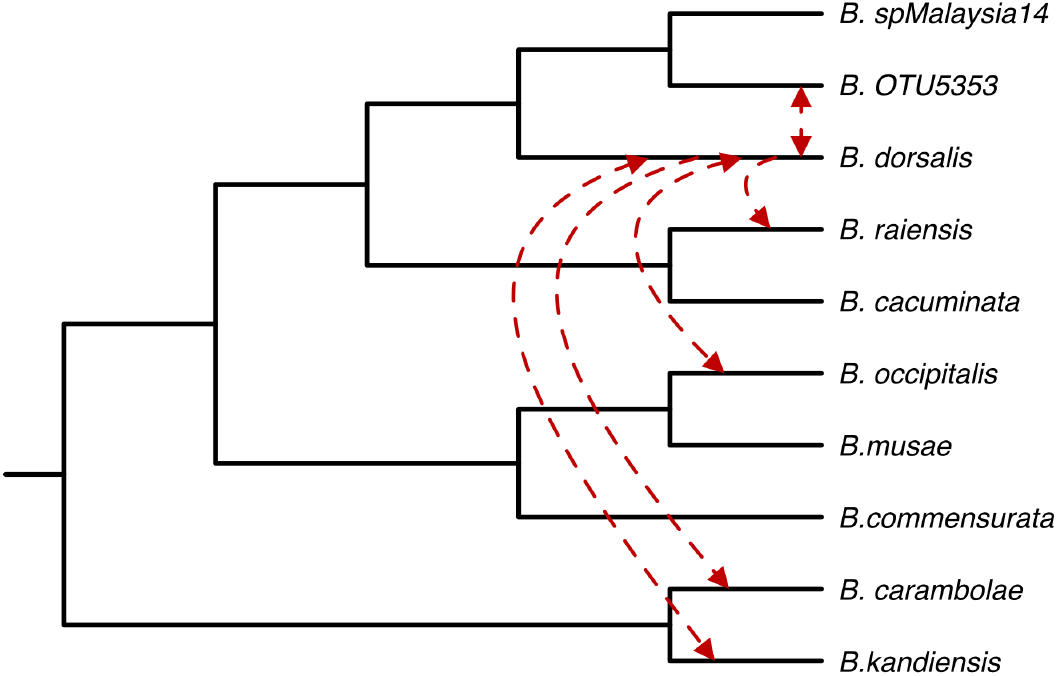
Cladogram with phylogenetic relationships of the *B. dorsalis* complex based on GBS genomic SNP data from San Jose et al. (in prep.). Red arrows indicate inferred horizontal COI transfer, presumably through mtDNA introgression, leading to false COI-based identifications.

### Bactrocera tryoni complex

The *Bactrocera tryoni* complex involves four Australasian cue lure attracted species that also encompass a morphological complex: *B. tryoni* (Froggatt 1897), *B. neohumeralis* (Hardy 1951), *B. melas* (Perkins & May 1949) and *B. aquilonis* (May 1965). Our results based on COI-5P3P are similar to those of COI-5P studies (Blacket et al. 2012; Jiang et al. 2014), which also could not distinguish the species. *Bactrocera tryoni*, the Queensland fruit fly, is a major agricultural pest in Australia and has invaded French Polynesia (Vargas et al. 2015) and it is intermingled with all other species in the complex. *Bactrocera tryoni* and *B. aquilonis* are disputably allopatric, with *B. aquilonis* restricted to the Northern Territory and *B. tryoni* found in eastern Australia. However, previous genetic studies based on microsatellites, like the COI results, have been unable to differentiate the two and their specific status has been much debated (Drew and Lambert 1986; Cameron et al. 2010). One hypothesis has been that they were originally two separate species, but that *B. tryoni* invaded the Northern Territory in the 1980’s and has since displaced the original *B. aquilonis*. *Bactrocera neohumeralis* can be found in the wetter areas of Queensland, where it is sympatric with *B. tryoni*. Its color patterns are slightly darker than those of *B. tryoni,* but the only discrete character that separates the two is the dark [*B. neohumeralis*] versus yellow [*B. tryoni*] color of the notopleural lobes. Microsatellite study supports that *B. neohumeralis* and *B. tryoni* are genetically distinct (Wang et al. 2003), and they differ in mating times (Morrow et al. 2000), but natural hybrids are known (Pike et al. 2003). *Bactrocera melas* is likely the most obscure member of the group, and it resembles a melanistic form of *B. tryoni* (Drew 1989). It can mostly be found in Southeast Queensland and is sympatric with *B. tryoni*. Taking ecology, distribution and nuDNA studies into account, it seems plausible that COI of the widespread *B. tryoni* introgressed into other members of the complex.

### Bactrocera mayi complex

The morphological *B. mayi* species complex involves three methyl-eugenol attracted flies in Australia, which we have sampled. We find that the *Bactrocera mayi* and *B. tenuifascia* COI clusters are mixed. Both species are considered allopatric with *B. tenuifascia* restricted to the Northern Territory and *B. mayi* to Queensland and surrounding areas, the morphological differences between them are minor and they may be candidates for synonymization.

### Bactrocera nigrotibialis complex

Three species of the morphological *Bactrocera nigrotibialis* complex are intermingled in the COI tree: *B. nigrotibialis* (Perkins 1938), *B. nigrifacia* Zhang Ji & Chen 2011 and *B. nigrifemoralis* White & Tsuruta 2001. *Bactrocera nigrotibialis* is a widespread species of economic concern that occurs from India to the Philippines and it can be found on most Indonesian islands, the other species have more limited distributions. Members of the *B. nigrotibialis* complex are typically only distinguished from each other by small differences in the black markings on the legs or face. None of the species in the *B. nigrotibialis* COI complex have received prior molecular attention with extensive sampling; it is possible that this is a case of incomplete lineage sorting, introgression, or inaccurate reference taxonomy.

### Bactrocera frauenfeldi complex

The Southeast-Asian *B. albistrigata*, Australasian *B. frauenfeldi* and Pacific *B. caledoniensis* are mixed in the COI tree. Similar results were found based on genomic data, but based on a small number of samples (Dupuis et al. 2018). Morphologically, they are separated based on the length of the yellow vittae on the scutum and a black mark on the scutellum, but they are considered strictly allopatric. *Bactrocera frauenfeldi* and *B. albistrigata* are damaging polyphagous agricultural pests, and further genomic or ecological studies could focus on the separation or syonymization of the species in this complex. The Pacific species *B. caledoniensis* and *B. trilineola* are further intermingled with a single specimen of *B. caledoniensis* falling in a clade with only *B. trilineola*. *Bactrocera trilineola* and *B. caledoniensis* are parapatric with overlapping distribution ranges in New Caledonia, after the relatively recent discovery of *B. trilineola* in the Loyalty Islands (Mille 2008), the two are morphologically highly similar.

## Discussion

### COI as an identification tool

Where early studies on the identification reliability of COI barcodes reported levels of non-monophyly as high as 26.5% across Arthropods (Funk and Omland 2003), these levels have gradually decreased as the operational error was reduced, to 16–18% across Arthropoda ten years later (Ross 2014). More recently, estimates based on a highly curated dataset of European moths and butterflies were at 12% non-monophyly, with half of those cases potentially attributable to operational error (Mutanen et al. 2016). A previous study on the COI reliability for Dacini fruit flies found 29% of the species non-identifiable (Jiang et al. 2014). For our dataset, we find 9.6- of the species overall non-monophyletic, and 21.4% of the pest species (Doorenweerd et al. 2018). Some of the non-monophyly may still be attributable to operational error such as inaccurate reference taxonomy. Of the 262 species covered in our dataset, 64 (24.4%) were singletons – species represented by a single individual. Although singletons can render other species non-monophyletic, they are always monophyletic by themselves and therefore inflate the reliability of COI identification statistics. With singletons excluded, the percentage of non-monophyly in our dataset is still moderate, at 12.6%. However, it has been shown that it may take hundreds if not thousands of representatives of a single species to adequately cover the intraspecific COI variation (Doorenweerd et al. 2019b; Phillips et al. 2019), and similar sample sizes may be required to detect introgression. For example, a previous study concluded that COI can reliably identify the Oriental fruit fly *Bactrocera dorsalis* (Blacket et al. 2012), but our study in conjunction with phylogenomic data (San Jose et al. in prep) finds that its COI is confused with five other species. To further improve our, and other, COI reference libraries would require two key efforts: increased sampling and phylogenomic data to support species-level identifications of the voucher specimens. As such, our current library for Dacini should be regarded as the most complete and curated COI dataset that is currently available, but requires further work to become a fully reliable reference for identifications.

### Validating reference libraries

One of the issues that has prevented the wide adoption of best practices of COI data (DeSalle and Goldstein 2019) is that a general set of statistics to indicate the reliability of a DNA barcode reference library has been lacking. Aside from including large tree figures, typical statistics reported in published papers include the presence, or more commonly, absence of a ‘DNA barcode gap’ – where the genetic variation between species is larger than within. This often includes a graph where all intra- and all interspecific pairwise-distance values of a dataset are plotted against each other (e.g. (Jiang et al. 2014; Zhang et al. 2017), but this approach suffers from two flaws. Firstly, the mean intraspecific variation increases as the taxonomic scope of the dataset widens, whereas for species identification the distance between sister species is the only relevant value (Dupuis et al. 2012). Secondly, non-distinct sequences (see M&M for definition) are typically not removed prior to estimations, and they distort the variation distribution within species. The Dmin_NN and Dmax statistics that we employed (see also (Ratnasingham and Hebert 2013), calculated after removing non-distinct sequences, more accurately indicate the reliability of identifications as datasets scale in sample-size and taxonomic breadth by focusing on nearest neighbor species. Ignoring ambiguous or missing data in the pairwise comparison of sequences further allows for more universal comparisons between sequence libraries, which now often report much inflated distance values and haplotype diversity. Although this should be regarded as only a first step, opening the discussion on how DNA barcode libraries can be validated and comparatively analyzed will be vital for their future potential.

### Haplotype ambiguity

Determining if two sequences are the same for pairwise distance estimates becomes increasingly difficult when ambiguous and/or missing data are involved (Johnson 2010). Ambiguous base-calls in DNA sequences can arise from multiple sources, and depending on the source, should be evaluated differently in down-stream analyses. For example, for nuclear DNA data in diploid organisms, it is possible to encounter heterozygous alleles, which are commonly encoded as ambiguous bases. In other cases, an ambiguous base represents a PCR or sequencing reaction artefact, or contamination, leading to a low level of confidence in the base call in the consensus sequence. For mitochondrial DNA, which is almost always maternally inherited and haploid (but see (Rubinoff et al. 2006), there is no reason for the natural presence of ambiguous bases in insects, except for rare cases of heteroplasmy (Rubinoff et al. 2006; Zhang et al. 2018), and ambiguous data therefore has no biological significance. As such, ambiguous bases and missing data should logically be ignored in pairwise comparisons. However, this is not how most pairwise comparison software currently functions, leading to bias in distance estimates (to the best of our knowledge; most pairwise comparison software is not open-source and does not specify methods). As compared to other methods, the PyCOIstats package that we developed and employed disregards ambiguous bases as potential differences, leading to reduced bias in the pairwise distance estimates that are used to evaluate the reliability and success rate of species-level identifications using COI. Future improvement of pairwise distance estimates may also involve different mutation models, although some studies have shown that uncorrected pairwise distance is most appropriate for COI data (Srivathsan and Meier 2011).

### Standardization versus optimization

The COI-5P segment had been touted to become the international standard for COI-based identification (Hebert et al. 2003a), at least for all animal life. The obvious advantage of standardizing the segment is that the data is more likely to be useable across different studies and it is a much more cost-efficient way to produce a universal reference library, instead of having to use multiple markers. However, standardization usually comes at a price, and the COI-5P fragment may not be the most informative or best at delimiting species for all Animalia. For Dacini, our results show that different segments of COI, provided they are larger than ~450 bp, have highly similar reliability for delimiting species, but work better for *Dacus* and *Zeugodacus* than for *Bactrocera*. However, an important yet often neglected aspect to consider with the selection of a DNA segment and the accompanying PCR primers is the potential for pseudogene amplification. At least two important pest species are known within Dacini where pseudogene amplification consistently hampers COI-5P identification (Blacket et al. 2012; Li et al. 2020). Amplifying a longer segment of COI in the PCR possibly avoids pseudogene issues, because nuclear copies of the functional gene quickly become fragmented by introns. However, this may not be feasible for samples with degraded and highly fragmented DNA, such as larvae in intercepted fruit or older museum specimens. In such cases, reliable amplification of the target segment may require non-universal species-specific primers to avoid pseudogenes (Blacket et al. 2012; Li et al. 2020). Species-specific protocols for molecular identification may thus be a necessity in addition to general, initial COI-based approach to reach a robust formal identification. The PacBio platform inherently provides an additional method to avoid pseudogenes: it generates a collection of independent single molecule reads for each sample, and this collection is then used to create the final consensus, revealing any off-target fragments or pseudogenes that are co-amplified clearly separated from other reads. The current drawback of the PacBio system is that it requires a large number of samples for a single run to be cost-effective. Including pseudogene avoidance through primer design, segment selection and sequencing platform in COI-targeting protocols will be crucial for a reliable molecular identification system.

## Acknowledgements

We thank the Sabah Biodiversity Centre for providing collection permits (ref. JKM/MBS.1000-2/3 JLD.3 (88)), And the help of Chi-Yeh Chien (Thai Royal Project Foundation), Thongsavanh Taipangnavong, Vornthalom Chanthavong (FAO insect pest management Laos), Lira Chea, Ajay Markanday (FAO insect pest management Cambodia), Johannes Ketellar (FAO insect pest management - Regional Office for Asia and the Pacific), Prabhat Kumar (Asian institute of Technology), Po-Yung Lai (University of Hawaii Mānoa CTAHR) and Shakil Ahmed Khan (Bangladesh Atomic Energy Commission) with permits and logistics in our international collecting trips, as well as all other institutions that provided collecting and export permits. We thank the following people for contributing specimens: J. Eiben, R. Putoa, A.Y. Kawahara, A. Ota, W.P. Haines, E.T. Vueti, N.-N. Zhang, G. Kang, D. Haymer, N. Howcroft, P.S. Aung, S. Cowan, B. Bhandari, T. Xian, E.J. Rodriguez, U. Freitas, S. Ouedrago, T. Stark, N. Pierce, S. Tam, F. Huda, Q.K. Le, T.T.T. Vu, Y.-C. Lin, J.B. Reil, T. Doi, S. Graham, K. Pohlman, Y. Xia, W.J. Lin, J.Y. Liang, K. Badji, A.L. Norrbom, G.J. Steck, J. Ismay, B. Ismay, J. Drouin, P. Waisen, E. Jang, C. Mille, K. Englberger and B. Bushe. We greatly appreciate help from Daniel Nitta with the molecular wet-lab work. We thank Carlos Lopez-Vaamonde for useful comments and suggestions on terminology. The technical support and advanced computing resources from University of Hawaii Information Technology Services – Cyberinfrastructure and the USDA SCINet scientific computing infrastructure are gratefully acknowledged. Funding for this project was provided by the United States Department of Agriculture (USDA) Plant Protection Act 7721. These funds were managed as interagency or cooperative agreements between USDA Animal and Plant Health Inspection Service and USDA-ARS (8130-0893-IA), the University of Hawaii’s College of Tropical Agriculture and Human Resources (8130-0565-CA) and the University of Idaho’s College of Agriculture and Life Sciences (8130-0665-CA). This material was made possible, in part, by a Cooperative Agreement from the United States Department of Agriculture’s Animal and Plant Health Inspection Service (APHIS). It may not necessarily express APHIS’ views. Additional funding was provided by the USDA Cooperative State Research, Education and Extension (CSREES) project HAW00942-H administered by the College of Tropical Agriculture and Human Resources, University of Hawaii. USDA is an equal opportunity employer. Mention of trade names or commercial products in this publication is solely for the purpose of providing specific information and does not imply recommendation or endorsement by the USDA.

